# Modeling Complex Animal Behavior with Latent State Inverse Reinforcement Learning

**DOI:** 10.1101/2024.11.13.623515

**Authors:** Aditi Jha, Victor Geadah, Jonathan W. Pillow

## Abstract

Understanding complex animal behavior is crucial for linking brain computation to observed actions. While recent research has shifted towards modeling behavior as a dynamic process, few approaches exist for modeling long-term, naturalistic behaviors such as navigation. We introduce discrete Dynamical Inverse Reinforcement Learning (dDIRL), a latent state-dependent paradigm for modeling complex animal behavior over extended periods. dDIRL models animal behavior as being driven by internal state-specific rewards, with Markovian transitions between the distinct internal states. Using expectation-maximization, we infer reward functions corresponding to each internal states and the transition probabilities between them, from observed behavior. We applied dDIRL to water-starved mice navigating a labyrinth, analyzing each animal individually. Our results reveal three distinct internal states sufficient to describe behavior, including a consistent water-seeking state occupied for less than half the time. We also identified two clusters of animals with different exploration patterns in the labyrinth. dDIRL offers a nuanced understanding of how internal states and their associated rewards shape observed behavior in complex environments, paving the way for deeper insights into the neural basis of naturalistic behavior.

## 1 Introduction

Understanding animal behavior is a central goal in neuroscience—it is crucial to link brain computations to observed behavior for a holistic understanding of the brain [17]. To this end, building computational models of animal behavior has proven to be an important step. These models provide interpretable descriptions of behavior, such as the various strategies animals employ during tasks [4, 3, 6] or modules that describe flexible behavior [7, 24]. These behavioral descriptions have also been linked to specific changes in neural activity, paving the way for a deeper understanding of how different neural circuits contribute to animal behavior [26, 6, 2, 21].

Historically, neuroscientists have characterized animal behavior as stereotyped in tasks such as binary decision-making [23, 9]. However, recent research has shifted towards modeling behavior as a dynamic, non-static process. For instance, binary decision-making behavior has been described as alternating between discrete strategies [4, 6], while social behavior in flies has been modeled through a similar hidden state-dependent process [7]. Furthermore, Wiltschko et al. modeled spontaneous rodent behavior as composed of behavioral “syllables” or motifs that animals switch between.

Navigation presents a complex yet naturalistic behavioral paradigm where animals make sequential decisions over time. While numerous computational models have been developed for decision-making settings [14, 10, 4], fewer modeling approaches exist for navigation-like behaviors [15]. In previous work [3], we developed a novel approach to model rodent navigation [20] as a dynamic process using inverse reinforcement learning. This approach characterized rodent behavior in terms of intrinsic time-varying reward functions, which provided an interpretable description of behavior. Notably, Ashwood et al. [3] found that these time-varying reward functions exhibited discrete switches over time, despite being modeled as continuous functions, echoing past work that modeled behavior as a combination of discrete states and syllables [25, 8, 4, 6]. However, [3] has two key limitations: (1) it requires pooling trajectories across multiple animals, thus limiting its applicability to long-term behavior modeling in individual animals, and (2) it lacks an explicit mechanism to model distinct behavioral states, despite evidence of their existence.

Motivated by these findings and limitations, we introduce discrete Dynamical Inverse Reinforcement Learning (dDIRL), a latent state-dependent paradigm for modeling long-term complex animal behavior. dDIRL posits that an animal’s actions are driven by internal state-specific rewards, with Markovian transitions between these internal states. Thus, it explicitly models the discrete nature of behavioral switches, and has fewer parameters than [3], making it more robust and applicable to individual animal behavior. Using an expectation-maximization algorithm, dDIRL allows us to infer both internal state-specific reward functions and transition probabilities between them, from observed behavior. For navigation tasks, we also implemented a smoothness prior on reward maps, ensuring that nearby locations have similar rewards.

We applied dDIRL to simulated gridworld environments and subsequently to water-starved mice navigating a labyrinth [20]. Unlike previous work [3], we analyzed each of the 10 animals individually, obtaining per-animal reward functions. Our results show that three distinct internal states sufficiently describe animal behavior in the labyrinth. One state consistently corresponded to a canonical water-seeking thirst state across all animals, which they typically occupy less than half of their time in the labyrinth. The remaining time was spent exploring other parts of the labyrinth. Intriguingly, we identified two clusters among the 10 animals: one group exhibited two reward maps capturing distinct exploration patterns, while the other showed one dominant exploration mode with occasional use of a third reward map.

Overall, this approach provides an interpretable description of animal behavior, enabling us to study both individual differences and similarities across animals performing the same task. By revealing the underlying reward structures and state transitions, dDIRL offers a nuanced understanding of animal decision-making processes in complex environments.

## 2 Discrete Dynamic Inverse Reinforcement Learning

### 2.1 Inverse Reinforcement Learning

We first describe the inverse reinforcement learning (IRL) problem. Let us consider a Markov Decision Process (MDP), ℳ = {𝒮, 𝒜, 𝒯, *r, γ*}, where 𝒮 is the state space, 𝒜 is the action space, 𝒯 : 𝒮 × 𝒮 × 𝒜 → [0, 1] represents the probability of transitioning between states when a certain action is taken, *r* : 𝒮 × 𝒜 → ℝ is the reward function, specifying the reward obtained by taking action *a* ∈ 𝒜 in state *s* ∈ 𝒮, and *γ* ∈ [0, 1] represents the discount factor. Inverse reinforcement learning [16, 1, 29, 5, 11, 27] aims to infer the unknown reward function *r*(*s, a*) when given access to {𝒮, 𝒜, 𝒯, *γ*} and *N* trajectories of agents navigating in this environment, 𝒟 = {*ζ*_1_, *ζ*_2_, …, *ζ*_*N*_}. Each trajectory is a sequence of state-action pairs, *ζ*_*i*_ = {(*s*_1_, *a*_1_), (*s*_2_, *a*_2_), …}.

### 2.2 dDIRL: Generative Model

Animal behavior has been shown to be non-stationary [4, 3], suggesting static reward functions inferred through inverse reinforcement learning insufficient for a comprehensive description of behavior. This was further validated by our earlier work [3], which while allowing static reward map as a potential solution, showed that the inferred weights of reward maps varied dynamically in time. We build and simplify in this direction by proposing a model where the animal transitions between distinct internal states over time.

We assume that at any time point *t* during a trajectory, the animal is in one of *K* distinct internal states, represented by *z*_*t*_ ∈ {1, …, *K*}. The reward function being optimized by the animal is dependent on its internal state. In order to avoid confusion between internal states *z*_*t*_ and the observed state *s*_*t*_, we refer to the latter as locations henceforth.

The reward corresponding to location *s* in the environment at time *t* in internal state *z*_*t*_ = *k* is given by:

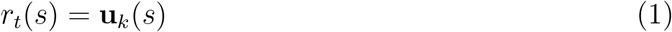

where **u**_*k*_ ∈ ℝ^|𝒮|^ is a static reward map (Fig. 1), capturing the animal’s reward function corresponding to internal state *k*. Each element, **u**_*k*_(*s*), reflects how rewarding the animal finds location *s* when in internal state *z*_*t*_ = *k*. We assume that the transitions between internal states are Markovian in nature, such that *z*_*t*+1_ depends on *z*_*t*_. Let *A* ∈ ℝ^*K*×*K*^ contain the probability of transitioning between internal states, such that *A*_*ij*_ = *P* (*z*_*t*+1_ = *j* | *z*_*t*_ = *i*). Furthermore, *ρ*_*z*_ ∈ Δ^*K*^ captures the probability of the initial internal state *z*_1_.

**Figure 1:**
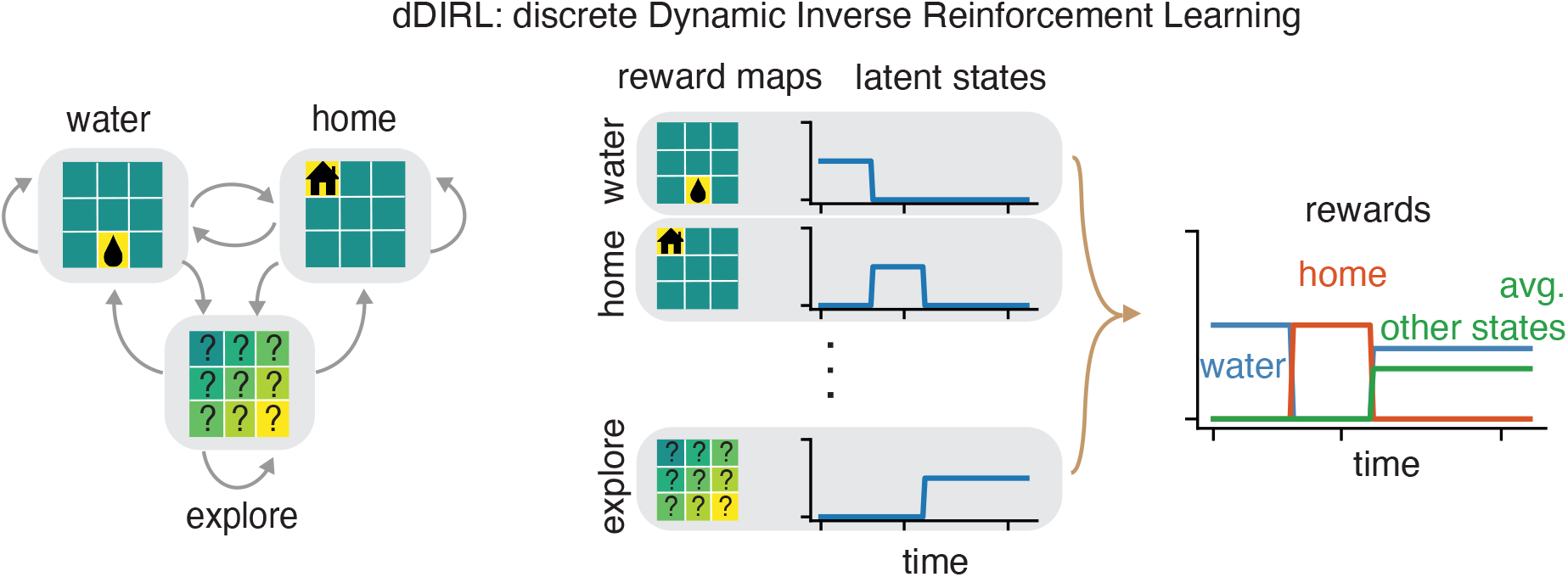
Schematic of discrete Dynamic Inverse Reinforcement Learning (dDIRL). (Left) Example reward functions corresponding to distinct internal states, arrows represent transitions between them. (Right) Example illustration of internal state-dependent rewards. Here, the water-seeking internal state is active initially, while the explore state is active in the end. The overall reward as a function of time thus reflects the reward map associated with the active internal state at any time point.

Next, following maximum likelihood inverse reinforcement learning [29, 28, 27], we assume that the animal optimizes a soft-max policy when navigating through its environment:

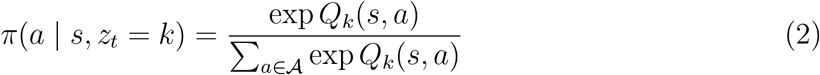

where *Q*_*k*_(*s, a*) is the soft-Q function when the animal is in internal state *k*. This represents the expected sum of future rewards starting from location *s* and action *a*, when the animal is in internal state *k*. Following Bellman equations, *Q*_*k*_(*s, a*) can be computed iteratively as follows:

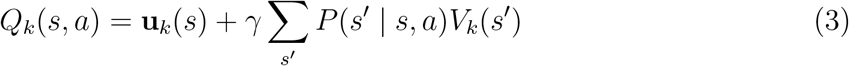

The first term, **u**_*k*_(*s*), is the reward value for the current location *s* (eq. 1) in the animal’s current internal state *k. V*_*k*_(*s*^′^) represents value function corresponding to location *s*^′^ in internal state *k*, capturing the expected sum of future rewards assuming the animal stays in internal state *k*.

Thus, the policy of the animal at any time point is dependent on its internal state *z*_*t*_, and is proportional to the sum of expected rewards starting from the current location *s*, and action *a*. As such, at time point *t*, the animal is likely to choose the action that maximizes its expectation of rewards starting at the current location-action pair {*s*_*t*_, *a*_*t*_}, and based on its current internal state *z*_*t*_. We think that this is a biologically plausible assumption, as the animal is presumably planning at any point conditional on its current internal state. We do not add an explicit temperature parameter in the policy, eq. 2, since our reward maps 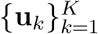 can take on any real value are not constrained to lie between {0, 1} as is the case in typical reinforcement learning setups.

Our model shares similarities with the work of Babes-Vroman et al., but extends their approach by allowing reward functions to change dynamically within trajectories, rather than only between them. Additionally, our work is closely related to the switching IRL model proposed by Surana and Srivastava [22], which was developed for surveillance applications. Our approach builds upon these foundations, adapting and expanding them to model complex animal behavior in naturalistic settings.

### 2.3 Inference Procedure

We learn the reward maps 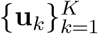, the internal state transition matrix *A*, and the initial state matrix *ρ*_*z*_, given access to the the observed trajectories 𝒟 using expectation-maximization (EM) [13]. Formally, our goal is to maximize the likelihood of the observed animal’s trajectories under our model, as follows:

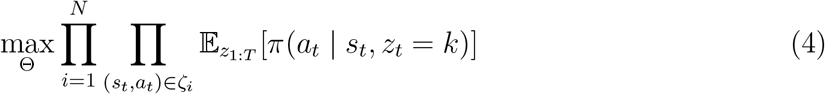

Where 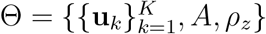. This resembles inference for Hidden Markov Models, where the observations of the model are location-action pairs.

In the E-step, we learn the probabilities of the internal states, *P* (*z*_*t*_ | Θ, 𝒟) and *P* (*z*_*t*_ = *j* | *z*_*t*−1_ = *k*, Θ, 𝒟) using the forward-backward algorithm, given the previously learned estimates of Θ (we initialize Θ randomly during the first iteration). Next, in the M-step, we learn an estimate of Θ given the above computed probabilities and the observed trajectories, by maximizing the complete data log-likelihood:

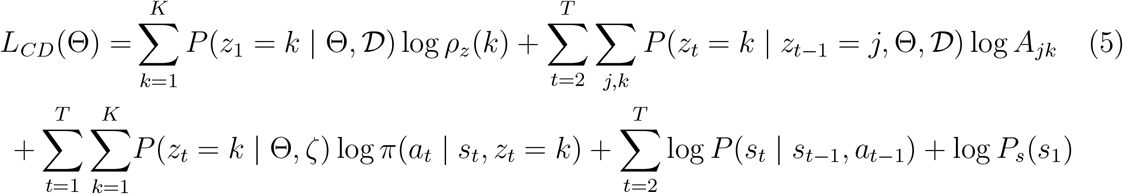

The Bayesian framework also allows us add priors to the model parameters, and maximize the log-posterior in the M-step. Specifically, we add a Dirichlet prior on each row of the transition matrix to encourage it to be sticky. For the *k*th row of the transition matrix, the Dirichlet parameters, ***α*** ∈ ℝ^*K*^, are such that *α*_*i*_ = 0 ∀*i* ≠ *k*, and *α*_*i*_ = *κ* ∀*i* = *k*. This results in the following closed-form updates for *ρ* and *A*:

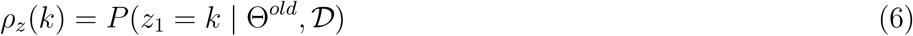

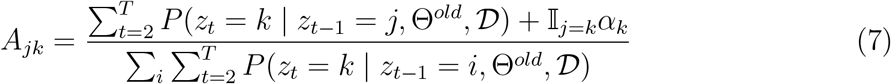

where the probabilities given Θ^*old*^ represent the posterior distributions over the internal latent states computed during the previous iteration of the E-step, and 𝕀_*j*=*k*_ is the indicator function.

We learn the reward maps using gradient ascent by maximizing the complete data log-likelihood, as they do not have a closed-form solution. We also add a smoothness prior so that proximity in space results in similar values of the reward values. Specifically, we add a graph-Laplace prior with an adjacency matrix designed to capture spatial location (depending on the specifics of the environment). This results in the following overall objective function:

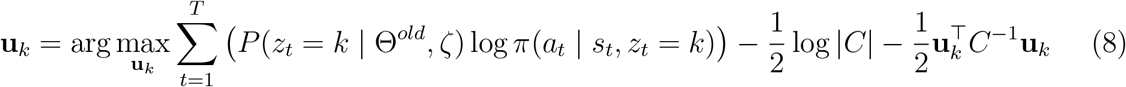

where *C*^−1^ = *D*^⊤^Σ^−1^*D*. Here *D* ∈ ℝ^|*S*|×|*S*|^ is the graph-laplacian where *D*_*ij*_ = −1, when *I* ≠ *j* and environment states *i* and *j* are adjacent, and is 0 when they are not adjacent. *D*_*ii*_ is set such that each row of *D* ultimately sums to 0. Σ^−1^ ∈ ℝ^|*S*|×|*S*|^ is a diagonal matrix with identical values, 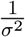, along the diagonals which tune the strength of the smoothness prior.

We learn the number of reward maps *K*, and the hyperparameters corresponding to the Dirichlet prior, *κ*, and the graph-Laplace prior, *σ*, using a held-out set of contiguous location-action pairs in our experiments.

## 3 Simulations in a gridworld environment

We illustrate our approach first in a gridworld environment. In a 5 × 5 gridworld environment, with 25 locations and 5 possible actions (left, right, up, down and stay), we generated two rewards maps (shown in Fig. 2B). One of these reward maps is highly rewarding at the top left location, gradually tapering off across the surrounding locations. The other map is highly rewarding at the bottom-center location, and similarly tapers off at nearby locations. These two reward maps correspond to two different internal states, where in one state the agent finds the top-left rewarding and in the other state it finds the bottom-center rewarding. The transition matrix, shown in Fig. 2C, is sticky with a high probability of staying in the current internal state.

**Figure 2:**
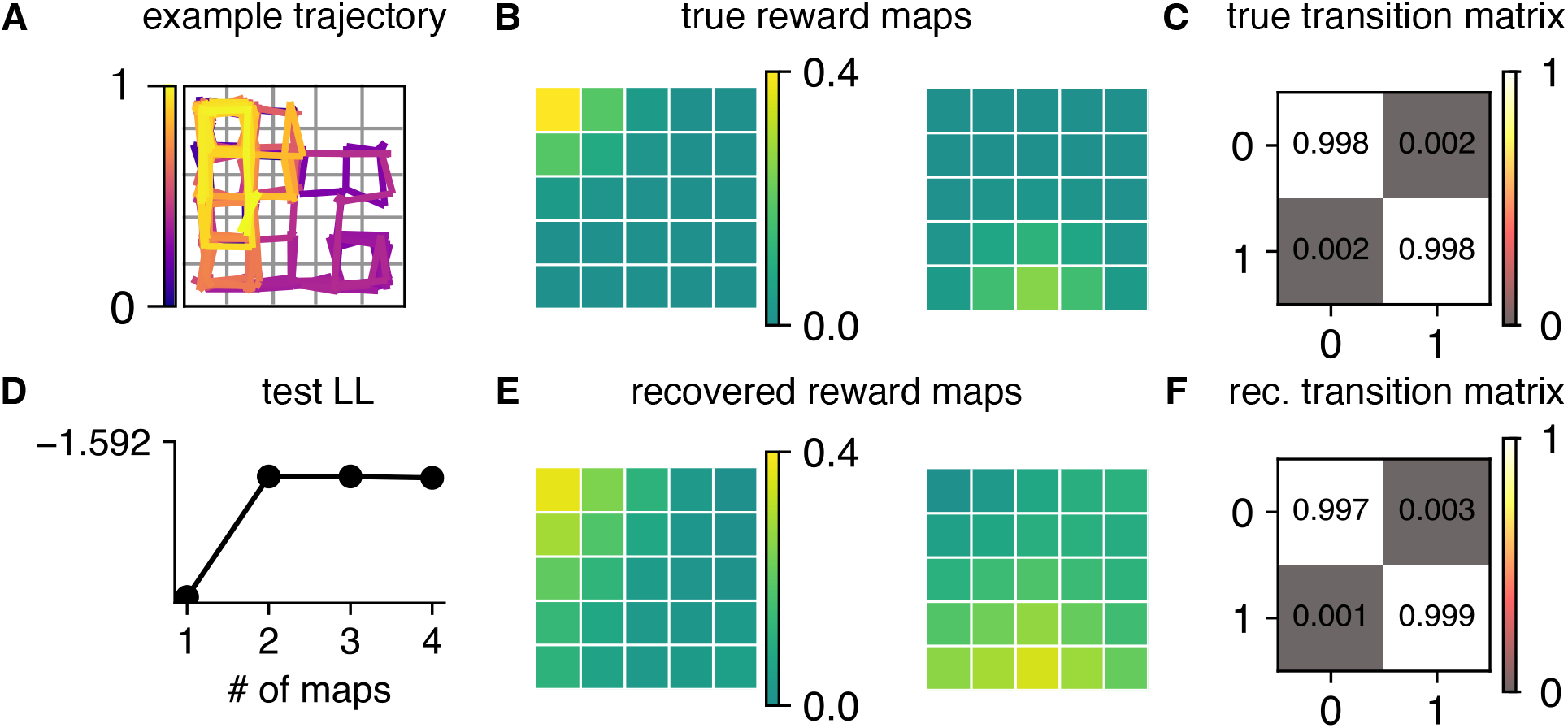
Simulations in a 5 × 5 gridworld environment. **A**. The first 500 time steps of an example trajectory in this environment, generated using parameters shown in **B & C. B**. True generative reward maps for two distinct internal states. The first map is highly rewarding near the top left location in the grid, while the second map is highly rewarding near the bottom center location. **C**. Transition matrix representing the probability of transitioning between the two internal states. **E**. Recovered reward maps using one trajectory of 5000 time steps. **F**. Recovered transition matrix. **D**. Test LL computed over a test trajectory at varying number of maps, it peaks at 2 maps which is the true number of states.

With this generative model, we generated a single trajectory of 5k time steps. We obtained the policy corresponding to each internal state using eq. 2. We then sampled internal states {*z*_*t*_} for the duration of the trajectory, and finally generated location-action pairs by rolling out the corresponding policy *π*(*a*_*t*_ | *s*_*t*_, *z*_*t*_) (the initial 500 time steps of this trajectory are shown in Fig. 2A).

We then used our EM-based inference algorithm to recover the reward maps and the transition matrix. We varied the number of maps in [1, 4], the Dirichlet prior *κ* ∈ [1, 1000], the graph Laplace prior *σ* ∈ [0.1, 1.0], and the learning rate to optimize the goal maps in [1*e*^−3^, 1*e*^−1^]. For each hyperparameter setting, we also initialized model parameters randomly with up to 10 different seeds, since EM is prone to getting stuck in local optima. We selected the best hyperparameters based on log-likelihood computed on a test trajectory of 1k time steps sampled from the same true generative model. We show in Fig. 2E & F that our inference scheme accurately recovers the reward maps (upto an additive constant^1^) as well as the transition matrix. Furthermore, Fig. 2D shows that the test log-likelihood peaks for 2 reward maps, which is the ground-truth in our generative model. Hence, these results validate our inference scheme to recover reward maps corresponding to distinct internal states and the transitions between them in a gridworld environment.

## 4 Inferring rewards for animals navigating a labyrinth

Next, we applied our approach to understand the behavior of mice navigating in a labyrinth environment. Rosenberg et al. recorded the trajectories of mice as they navigated through a labyrinth environment over the course of hours in the dark (Fig. 3). Ten of these mice were water-restricted. The labyrinth was structured in the form of a binary tree with 127 nodes, and one node provided water to these animals. In previous work [3], we obtained a reward functions across all trajectories of these 10 animals, and observed that it changed over time. Here, we applied dDIRL to the trajectories of each of these 10 mice separately.

**Figure 3:**
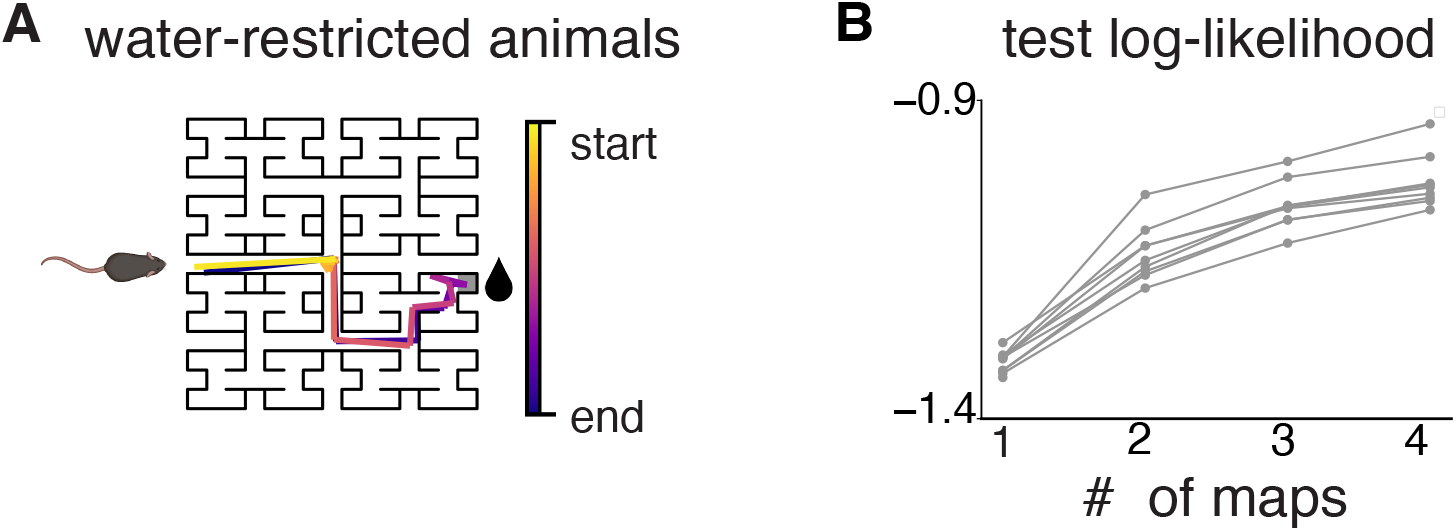
dDIRL applied to water-restricted mice performing the labyrinth task. **A**. Water-restricted mice navigated the labyrinth environment in the dark [20]. The labyrinth had 127 nodes, with only one node providing water (marked with a water drop). **B**. We applied our internal state-based IRL approach to trajectories of 10 such mice, and found that test log-likelihood (on held out time steps) typically saturates at 3 maps.

As previously mentioned, the labyrinth environment consists of 127 nodes, which form the set of locations within our environment. With the exception of terminal nodes, the animal could go left, right, backwards or stay at each node. At terminal nodes, it could only stay or go back. For each animal, we excluded the initial 20 minutes of data, as during this phase, the animal is learning to navigate the labyrinth and is unaware of the structure of the environment [20]. We sampled location-action pairs for each animal at a frequency of 1Hz resulting in 6, 600 − 19, 000 state-action pairs per animal, with an average of 15, 000 pairs per animal.

We fit internal state-based IRL to each of the 10 animals separately, while varying the number of maps between 1 − 4. We also varied the Dirichlet prior, *κ* ∈ [1, 1000], the graph Laplace prior *σ* ∈ [2^1^, 2^8^], the learning rate to optimize goal maps in [1*e*^−3^, 1*e*^−1^], as well as the discount factor *γ* ∈ [0.6, 0.9]. To infer these hyperparameters, we held-out 20% of contiguous location-action pairs from each animal’s trajectory.

### 4.1 Three reward maps underlie navigation behavior, with individual variability across animals

As shown in Fig. 3, we found that across all 10 animals, typically the test log-likelihood saturates around 3 maps. This is an exciting finding, and in line with existing work that a small number of discrete modules well explain animal behavior [4, 6, 24]. Upon examining the reward maps corresponding to individual animals, we consistently found a water-seeking reward map across all animals. This is a crucial validation of our approach, as we expect water-seeking animals to find the water port rewarding (Fig.4**A & B**, leftmost plot). Moreover, each animal had a sticky transition matrix, so that the probability of staying in the current internal state was high (Fig.4**A & B** rightmost plots).

Subsequently, for each animal, we determined the most probable internal state at each time step and examined the proportion of instances in which the animal utilized a particular reward map. We then averaged the computed proportions for the water-seeking map, and separately averaged the proportions for the remaining two maps after sorting them in order of utility. Firstly, we find that all animal spend about 30 − 40% of time in their water-seeking internal state (Fig. 4C, blue bar). The remaining time is spent exploring different parts of the labyrinth. Intriguingly, based on the remaining two reward maps that capture exploration behavior, the animals could be clustered into two groups containing 5 animals each. While one group of animals utilized all three inferred maps in equal proportions, the other group predominantly utilized two of the three maps, with only a small fraction associated with the third map (Fig. 4C).

**Figure 4:**
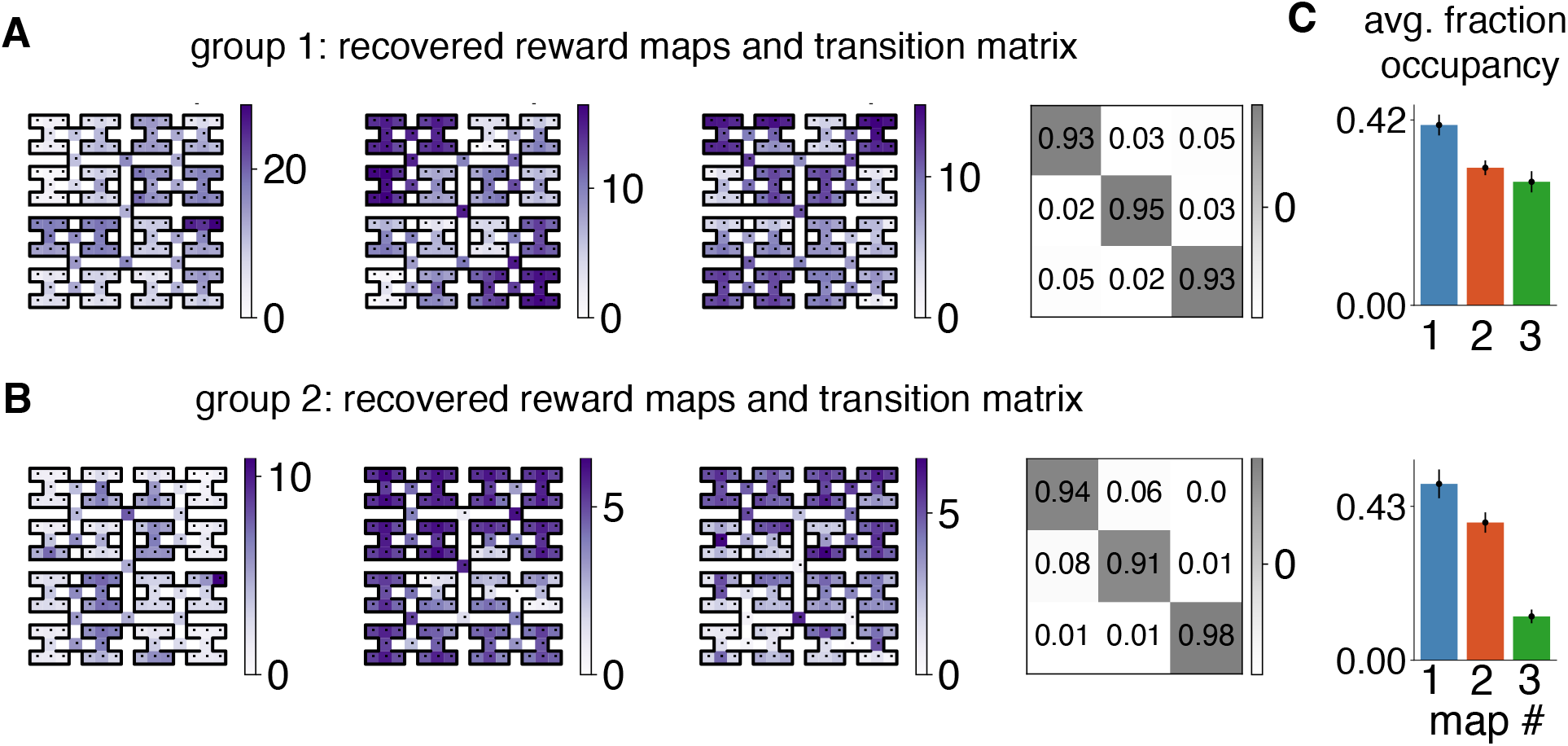
Our method reveals two groups of animals based on their exploration behavior. **A**. Recovered parameters for an example animal in group 1. This group of animals had a water-seeking map, and two different exploration maps capturing distinct modes of exploration behavior. *(From left to right)* Water-seeking map, followed by two exploration maps, and the recovered transition matrix. **B**. Recovered parameters for an example animal in group 2. This group of animals also had a water-seeking map, and only one dominant exploration map, the third map was only occasionally used. *(From left to right)* Water-seeking map, dominant exploration map, occasional exploration map, and the recovered transition matrix. **C**. Average fraction of times each map was used across all animals. Blue bar represents the water-seeking map across all animals. Orange and green refer to the two exploration maps recovered per animal, sorted in their order of utility. (*Top*) Animals in group 1. (*Bottom*) Animals in group 2. Errorbars show 95% confidence interval across animals.

We show the recovered reward maps for an example animal in Fig. 4A (top), where the map in the center is rewarding at the top-left and bottom-right corners of the labyrinth, and the right map is most rewarding towards the top fringes of the labyrinth. While there is individual variability across animals in terms of the locations they find rewarding, we found that all animals in this group could be consistently described by two maps capturing exploration behavior along with a water-seeking map.

The second group of 5 animals also had interesting similarities—they spent most of their time exploring based on one dominant exploration map. They only occasionally use a third map, which for some of these animals was most rewarding at junction nodes in the labyrinth, presumably capturing time steps when the animal is deciding which part of the labyrinth to go to next. In Fig. 4A (bottom), we show recovered maps for an animal in this group, where the the map in the center captures the animal’s predominant exploration behavior—the animal found the top half of the labyrinth to be most rewarding. The map in the right is most rewarding at several junction nodes, and was only occasionally used.

Finally, we can use the learned parameters to infer the underlying internal state of the animal at any point. This can allow us to divide animal behavior into discrete segments, and can further be correlated with neural activity in future work. We show the first few time steps of trajectories of two animals in Fig. 5A, colored by the internal state of the animal alongside a time series of their internal states. While the animal in group 1 (Fig. 5 top) switches between the 3 distinct internal states, an example animal from group 2 (Fig. 5 bottom) is typically in one of two internal states (using either the water-seeking map or the dominant exploration map).

**Figure 5:**
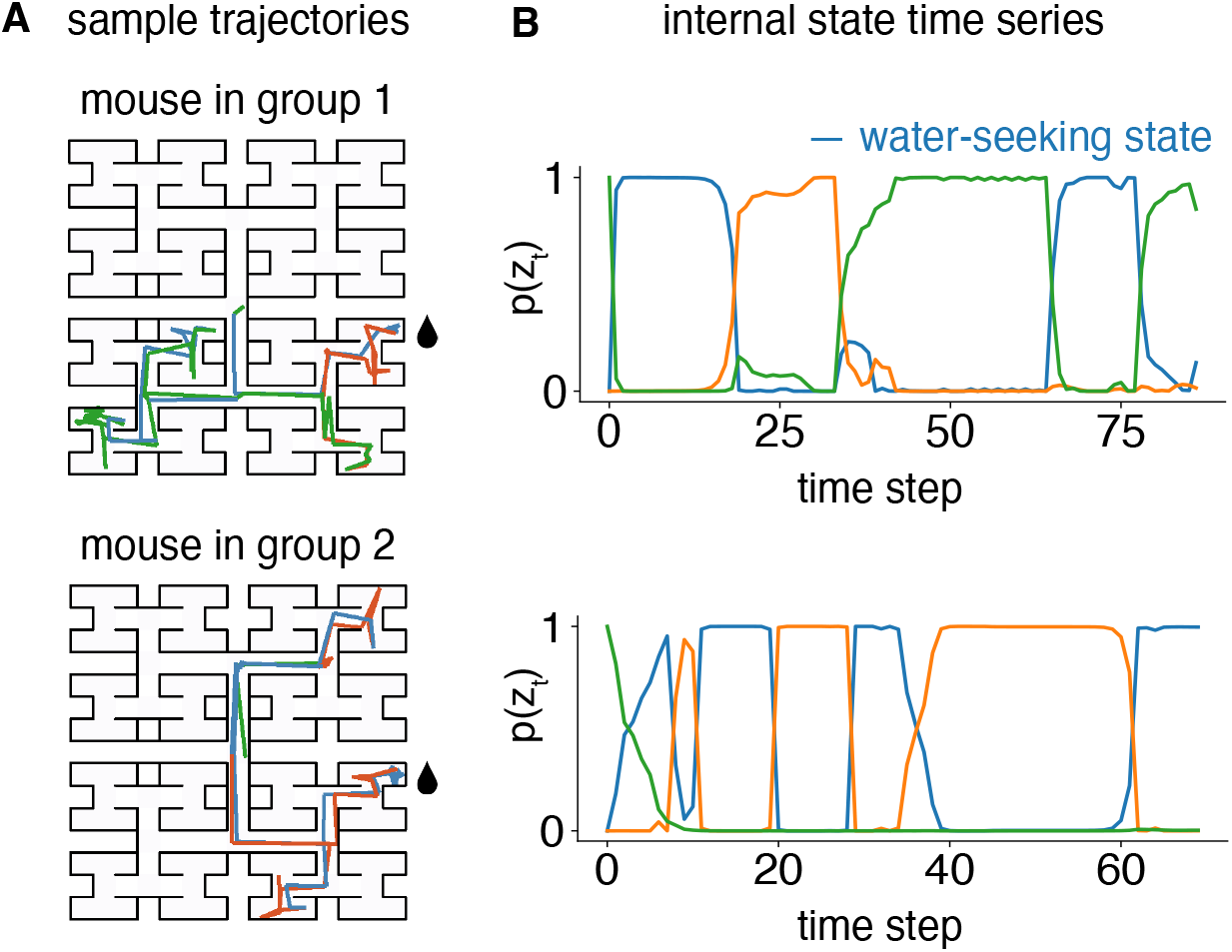
Inferred internal state along segments of animal trajectories. **A**. *(Top)* First 90 time steps of the trajectory of an animal from group 1, colored by its internal state. Blue represents the water-seeking state. *(Bottom)* First 70 time steps of the trajectory of an animal from group 2 colored by its internal state. We can see that animal spends time at the water port, shown by the jittered blue lines at the water port. **B**. The probability of each internal state plotted as a function of time for the two segments shown in **A**. *(Top)* Inferred internal states for the mouse from group 1. (*Bottom*) Same for group 2.

In summary, our results highlight the significance of comprehending behavior at an individual animal level. While aggregating data across animals provides insights into overall behavior trends, each animal exhibits unique characteristics that can only be revealed through independent analysis of their trajectories.

## 5 Discussion

In this work, we developed an latent state-based inverse reinforcement learning approach to study animal behavior during navigation. We infer the intrinsic reward functions motivating animal behavior, and parameterize these reward functions to depend on the animal’s internal cognitive state. We validated our framework in a simulated gridworld environment, and then applied it to study animal behavior in a labyrinth environment [20]. For each animal, we had access to one long trajectory spanning hours. Intriguingly, we found that for all 10 animals, three reward functions described their behavior well. One of these maps was a water-seeking map, as these animals are water-starved. However, they spent less than half their time seeking water, and spent the rest of their time exploring the labyrinth. While we observed individual variabilities across animals, overall animals could be grouped into two cohorts. One cohort had 2 distinct modes of exploration captured by two distinct exploration maps, the other cohort had one dominant mode of exploration and only occasionally used a third map.

The discovery of reward maps on a per-animal is particularly exciting as this allows us to study individual-to-individual variability in animals. Future work can link this variability to the learning phase of these animals [19], in order to understand the normative cause behind this variability. Furthermore, this approach allows us to segment hours of animal behavior based on the animal’s internal state. This can then be correlated to neural activity, for example, to understand which neural computations and regions are involved in exploration and how animals switch between these modes [2]. Finally, while we assumed switches between the animal’s internal states to depend only on its previous internal state, the location / environmental state of the animal can also affect its internal state. In future work, we thus hope to include environmental state as a covariate in the animal’s internal state transition matrix [7].

While we applied our approach to rodent behavior in a labyrinth, it can be applied across a variety of settings to study flexible animal behavior, such as goal-directed navigation in flies [12]. With the growing interest in understanding animal behavior [4, 26, 18], our paradigm adds a new perspective and approach to modeling animal behavior where we understand behavior from the perspective of the animal’s goals and action, along with its internal state.

## 6 Acknowledgements

We thank Zoe Ashwood and Scott Linderman for fruitfuil discussions throughout this project. AJ was supported by a Google PhD fellowship. VG was supported by doctoral scholarships from the Natural Sciences and Engineering Research Council of Canada (NSERC) and the Fonds de recherche du Québec – Nature et technologies (FRQNT). JWP was supported by grants from the Simons Collaboration on the Global Brain (SCGB AWD543027), the NIH BRAIN initiative (9R01DA056404-04), an NIH R01 (1R01EY033064), and a U19 NIH-NINDS BRAIN Initiative Award (U19NS104648).

## Footnotes

1 The reward function is non-identifiable as adding a constant to **u**_*k*_ does not change the softmax policy. Thus, we post-process the reward functions so that they have a minimum value of 0.

## Notes

### Competing Interest Statement

The authors have declared no competing interest.

